# Contrasting patterns of local adaptation and climate resilience across forest management regimes in Norway spruce (*Picea abies):* implications for reforestation practices under climate change

**DOI:** 10.1101/2025.06.11.659044

**Authors:** Helena Eklöf, Carolina Bernhardsson, Pär K. Ingvarsson

## Abstract

This study investigates the contrasting patterns of genetic diversity and local adaptation between old-growth and recently planted Norway spruce (*Picea abies*) stands in northern Sweden. We assess neutral genetic variation, adaptive genetic differentiation, and the potential for future adaptation using samples collected from old-growth forests and recently planted stands. Our results reveal no significant differences in overall genetic diversity between natural and planted populations, indicating that current forest management practices have not substantially reduced genetic variation. However, analyses of adaptive variation demonstrate stronger signatures of local adaptation in old-growth populations, with clear correlations between genetic and environmental distances. In contrast, planted stands show weaker adaptive signals and are at greater risk of non-adaptiveness under future climate scenarios. These findings highlight the importance of conserving and promoting adaptive genetic variation within forest management to ensure resilience against ongoing climate change, while also recognising that current practices preserve much of the neutral genetic diversity necessary for long-term forest health.

## Introduction

Forest management aims to increase ecosystem benefits and services over those expected from unmanaged forests (Andersson *et al*., 2017). Management practices have been shown to affect forest diversity, from the genetic level to populations and ecosystems (Lefèvre, 2004; Aravanopoulos, 2018). Ongoing climate change is also expected to impact forest ecosystems, and even if most forests have extensive adaptive capabilities, adaptive forestry practices are needed for managed forests to respond to climate-driven changes (Aitken *et al*., 2008; Alberto *et al*., 2013; Aravanopoulos, 2018; Fady *et al*., 2020). One of the most important factors to consider when managing forests for climate change is the preservation of genetic diversity, as high levels of genetic diversity are essential for preserving the adaptive capacity of forest populations (Aitken *et al*., 2008; Lefèvre *et al*., 2013). Forest management and silviculture can significantly impact the genotypic and phenotypic variation in forests, for example, by selecting trees for felling or choosing strategies for reforesting following harvests. Several methods for reforesting are in use in contemporary commercial forestry, including leaving natural seed trees, direct sowing of seeds or planting pre-established seedlings (Nilsson *et al*., 2010; Aravanopoulos, 2018). Planting genetically improved seedlings obtained from nurseries is the prevailing method in many parts of the world, as this speeds up reforestation and allows for more even stands while reducing the need for thinning. For example, 84% of all reforestation in Sweden is achieved through seedling plantations, most of which are obtained from seed orchards made up of genetically improved material. Natural regeneration makes up an additional 10%, and direct seeding and no measures taken (where the regeneration methods are unknown) make up the remaining fraction of new stand establishments in Sweden (Black-Samuelsson *et al*., 2020). Although large-scale reforestation has been ongoing since the mid-20th century in many parts of the world, it is unclear to what extent such practices, especially in combination with genetically improved material from existing breeding programs, alter genetic diversity in planted forest stands. Using improved seeds originating from a seed orchard consisting of a relatively small number of unique, genetically improved trees could reduce the effective population size of newly planted stands and hence limit genetic diversity (Savolainen & Kärkkäinen, 1992; Ingvarsson & Dahlberg, 2019).

Several studies have compared genetic diversity between managed and natural stands of different conifer species to assess these possible concerns. Bergmann and Ruetz (1991) used isozyme gene markers and found no significant difference in gene diversity between forest samples and seed orchard clones in a forest district in Germany. They did, however, see a difference in the average heterozygosity, suggesting slight differences in genetic diversity between the two types of samples. Maghuly et al. (2006) performed a study in Austria using microsatellites derived from both mitochondrial DNA, chloroplast DNA and nuclear DNA to compare two age groups (6-10 years and 70-100 years) from three different subpopulations (derived from three different elevations). The nuclear SSR markers showed a slightly higher level of genetic variation within populations of both age groups than genetic differentiation among subpopulations. Similarly, Ruņģis et al. (2019) used 11 SSR markers to compare naturally regenerated Norway spruce (*Picea abies*) populations to progenies derived from two different seed orchards in Latvia. Ruņģis et al. (2019) found that the total and effective number of alleles, average number of alleles, average gene diversity and average allelic richness were higher in the naturally regenerated forests compared to the seed orchards. Genetic diversity indicators were similar in all populations, and genetic diversity in the progeny from seed orchards was comparable to levels from naturally regenerated forests. Finally, in Scots pine (*Pinus sylvestris*), nuclear and chloroplast-derived microsatellites were used to compare genetic diversity in natural forests, seed-tree-generated forests and stands planted using genetically improved seedlings originating from three regions in Sweden (Garcia-Gíl *et al*., 2015). The results showed that reforestation methods did not affect nuclear or chloroplast genetic diversity. However, the number of effective alleles and total gene diversity in chloroplast markers were significantly higher in stands from seed trees compared to natural forests and seedling planting stands (Garcia-Gíl *et al*., 2015).

Previous studies assessing genetic diversity in natural and managed stands of forest trees (Bergmann & Ruetz, 1991; Maghuly *et al*., 2006; Garcia-Gíl *et al*., 2015; Sønstebø *et al*., 2018; Ruņģis *et al*., 2019) have been based on either low-resolution markers (e.g. allozymes) or have relied on a small number of high-resolution markers (e.g. microsatellites) that have limited the statistical power for detecting systematic differences in genetic diversity or differentiation. Such markers are also generally not considered directly involved in controlling adaptive traits and, therefore, convey little information on the present and future adaptive potential of different forest stands. Earlier studies have shown that patterns of adaptive variation are often strikingly different from those seen in neutral genetic variation (Le Corre & Kremer, 2003; Savolainen *et al*., 2013). Natural selection favours alleles that increase adaptation to local growing conditions, often resulting in increased genetic differentiation among populations for traits that contribute to local adaptation (Savolainen *et al*., 2007, 2013). In most forest trees, such local adaptation persists despite high levels of gene flow that continuously introduces potentially maladaptive genetic variation (Savolainen *et al*., 2007; Kremer *et al*., 2012). Most neutral markers discussed in the preceding sections usually display low genetic differentiation among forest tree populations, consistent with the extensive and homogenising effects of gene flow. Genetic differentiation at loci underlying adaptive traits is instead expected to show enhanced genetic differentiation due to diversifying effects of local selection and possibly reduced gene flow (Savolainen *et al*., 2007, 2013). However, in many cases, adaptive loci nevertheless display weak differentiation but instead show marked covariances in allele frequencies across populations rather than detectable changes at individual loci (Latta, 1998; Le Corre & Kremer, 2003; Kremer & Le Corre, 2012). Modern forest tree breeding programs typically include climate adaptation as a primary breeding objective (Holliday *et al*., 2017; Cortés *et al*., 2020; Isabel *et al*., 2020), however, to date, few studies have addressed how current forest management practices alter adaptive genetic variation and local adaptation *in situ*.

Norway spruce (*Picea abies* (L.) Karst.) is one of the most important conifer species in Europe, with a distribution range extending from the west coast of Norway to mainland Russia in northern Europe and across the Alps, the Carpathians and the Balkans in central Europe (Giesecke & Bennett, 2004). In 2013, there were around 3.5 billion cubic meters of standing forest on productive forest land in Sweden, with 41% of Norway spruce and 39.1% of Scots pine (Black-Samuelsson *et al*., 2020). Norway spruce is characterised by large population sizes and low mutation rates that, together with being an outcrossing and wind-pollinated species, lead to a high genetic diversity within and a low differentiation among populations (Burczyk *et al*., 2004). Breeding of Norway spruce in Sweden was initiated in the 1950s by selecting plus trees (i.e. trees with phenotypically desirable qualities such as straight trunks, free from damage and of high quality) from natural forests. Today, breeding populations consist of trees of both Swedish and foreign origin, which are included to improve genetic gain and are extensively evaluated using field testing. Material from the breeding populations has been used to establish commercial seed orchards designed to maximise genetic gain in value and volume production (Lindgren *et al*., 2007). The recommended optimal number of unique genotypes included in a seed orchard, to maximise the estimated benefit, has been estimated to be 16 (Lindgren & Prescher, 2005), assuming unrelated parent trees with normal fertility variation and that any pollen contamination is derived from unrelated sources (Lindgren & Prescher, 2005).

The genetic diversity of Norway spruce seed orchard crops has been investigated by Sønstebø et al. (2018) using 11 microsatellites. They compared two seed orchards, consisting of 25 and 60 parent trees, with seed crops collected from semi-natural or unmanaged forests from the same regions (Sønstebø *et al*., 2018). They showed that genetic diversity in orchard seed crops was slightly lower than in seed crops collected from semi-natural or unmanaged forests (Sønstebø *et al*., 2018). Similarly, nine breeding populations from northern and central Sweden were examined using 15 simple sequence repeat (SSR) markers, and the results showed high genetic diversity within populations, low genetic differentiation between populations and low inbreeding and relatedness levels (Androsiuk *et al*., 2013). Finally, Verbhylaité et al. (2017) evaluated genetic diversity in self-regenerated populations of Norway spruce and Scots pine in Lithuania. They observed high genetic diversity and low genetic differentiation between maternal trees and regenerating seedlings, suggesting that self-regenerating populations can provide sufficient genetic diversity to ensure ecologically and evolutionarily sound stands (Verbylaitė *et al*., 2017).

In this study, we assess genetic diversity in a selection of old-growth stands of Norway spruce from northern Sweden that have not been subjected to logging for at least 150 years. We contrast the natural, old-growth stands with similar data obtained from recently planted stands (<25 years) located near the old-growth populations. Our goal is to assess how patterns of genetic diversity differ between the two stand types and to identify genetic variation associated with local climate. We also evaluate the relative importance of isolation by distance and isolation by environment (Wang & Bradburd, 2014) for the different types of stands. Finally, we use climate modelling data to assess the risk of maladaptation of the different types of stands to projected climate changes predicted for the end of the 21st century.

## Method

### Sampling

We cooperated with Länsstyrelsen Västerbotten to identify and obtain coordinates for old-growth forest stands in the Västerbotten and Västernorrland counties of northern Sweden (Figure 1, Supplementary Table 1). We selected 15 stands without recorded logging for at least 150 years. These stands ranged from the Baltic coast in the east to the Scandinavian mountains in the west, capturing the east-to-west distribution of Norway spruce (*Picea abies*) in northern Sweden (Figure 1, Supplementary Table 1). Near each old-growth stand, we identified two newly planted forest stands (planted <25 years ago). We obtained stand coordinates, information on the origin of planting material and date of replanting, where available, from two forest companies active in the region, SCA Skog AB (11 stands) and Sveaskog (19 stands). For the Sveaskog stands, we obtained the actual year of replanting, while for the SCA stands, we obtained the year when clear-cutting had been performed (usually one or two years before replanting).

**Figure 1.**
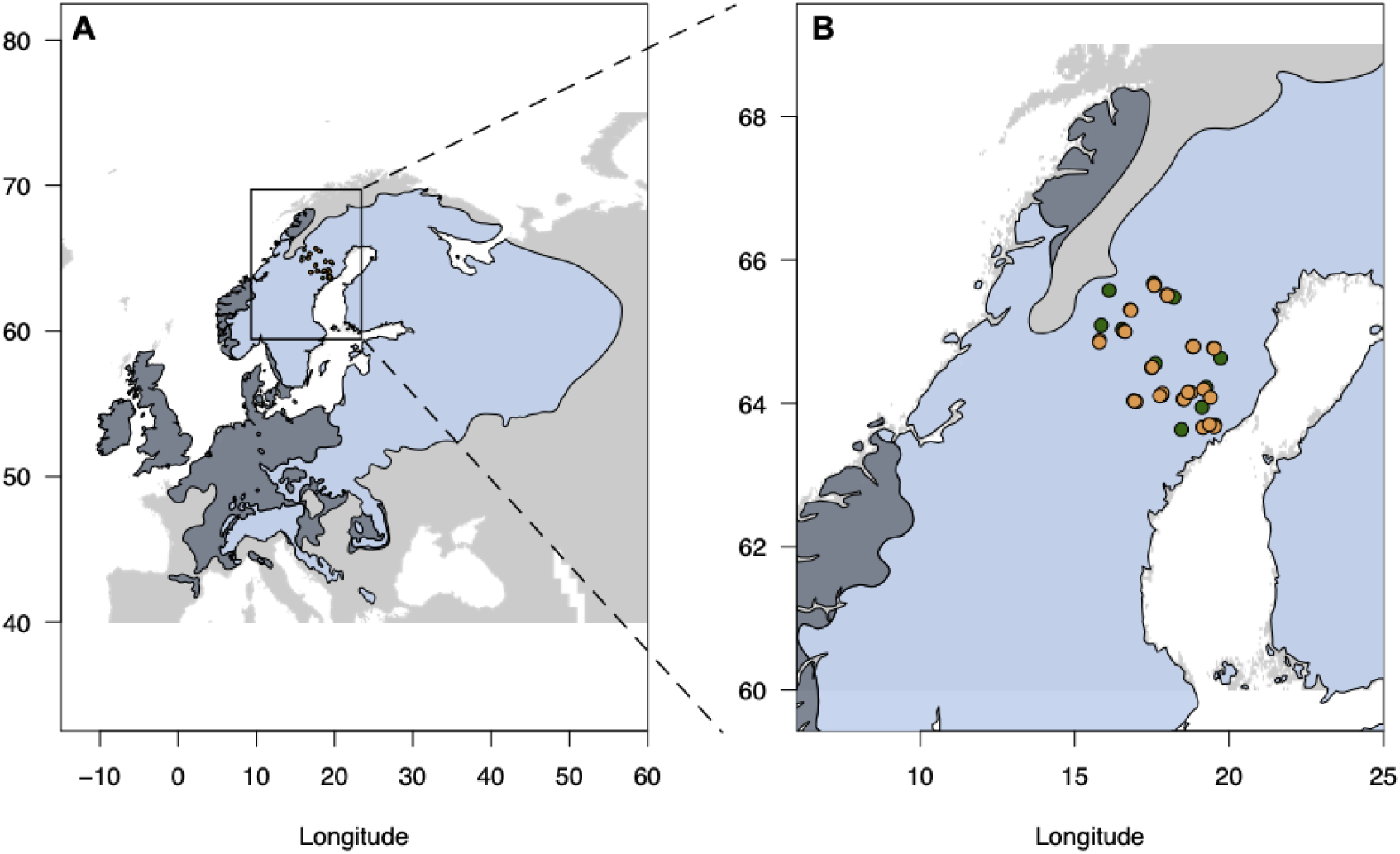
A) The natural (light blue) and introduced (dark slate) distribution range of *P. abies* in Europe. B) Close-up of northern Sweden with the sample sites highlighted in green for old and tan for planted populations.

In June 2017, we visited all 15 old and 30 young re-planted stands and sampled newly flushed buds from 50 random trees using transect sampling to obtain material from 2250 trees. All samples were kept on ice while sampling in the field and then brought back to the lab and stored at −80 °C until DNA extraction. From the 50 samples per stand, we randomly selected 30 samples for a total sample size of 1350 trees and used these for DNA extraction.

### DNA extraction, sequencing and SNP calling

Population genetic analyses rely on accurately estimating allele frequencies from population samples. Sequencing pooled samples of individuals (Pool-Seq) have been shown to provide more accurate allele frequency estimation than sequencing individuals because more individuals can be assessed with a Pool-Seq approach and usually at only a fraction of the cost compared to sequencing separate individuals (Schlötterer *et al*., 2014). To be able to genotype as many trees and stands as possible, we chose to adopt a Pool-Seq strategy for genotyping our samples.

DNA was extracted from all samples using an Omega Bio-tek E-Z 96 plant kit (OMEGA Bio-Tek), and DNA concentrations were measured on a Qubit (ThermoFisher Scientific). High-concentration samples were diluted, and 30 samples from each stand were pooled in equimolar concentrations by using 200 ng of DNA from each sample. Each pool was divided into eight tubes (750 ng DNA per tube) to keep the reaction volume low. The method we used for GBS library preparation is described in detail by Pan et al. (2015), with some minor changes described below.

DNA digestion and ligation were performed in 50 ⎧l reagent systems using the restriction enzyme *Pst*I (New England BioLab, Woburn, MA, USA). Adaptors were ligated to each of the 45 DNA stand pools using five unique barcodes. Five stand pools with unique barcodes were further pooled to create a super-pool. The nine super-pools (5 stand pools per super-pool) were purified using a QIAquick PCR purification Kit (Qiagen, Hilden, Germany) and DNA concentrations were measured on a Qubit (ThermoFisher Scientific). All nine super-pools were amplified using a PCR step with 12 50 ⎧l reagent systems for each pool and then purified. Size selection was made with the E-gel Size-Select II pre-cast gel (ThermoFisher Scientific), using a total of 100 ⎧l sample from each super-pool. Targeting fragments in the range 350-450 bp (accounting for 125–132 bp barcodes and sequencing adapters), the gel was run for approximately 20 minutes before the desired fragment size was excised from the gel. The gel was cut, DNA was extracted using a QIAquick Gel Extraction Kit (Qiagen, Hilden, Germany), resulting in nine unique super-pool libraries. Each library was quantified on a Qubit (ThermoFisher Scientific), and pair-end sequencing (2×150 bp) was performed on an Illumina HiSeqX by Novogene Europe. Each super-pool was sequenced individually on one HiSeq X lane, yielding > 120 Gbp raw sequencing data per super-pool.

The raw sequencing data were quality checked using FastQC v0.11.8 (https://www.bioinformatics.babraham.ac.uk/projects/fastqc/), and sequencing adaptors were trimmed using Trimmomatic v0.36 (Bolger *et al*., 2014). The sequence data from each super-pool library were demultiplexed using the `process_radtags` routine from Stacks v2.2 (Catchen *et al*., 2011) to extract reads for the individual stand pools. All sequencing reads were mapped against the *P. abies* v1.0 genome (Nystedt *et al*., 2013) using BWA-MEM with default parameters (Li, 2013). For SNP calling, we used `samtools` v1.14 (Li *et al*., 2009) to create an `mpileup` file from all 45 individual BAM files and SNPs were then called from this file using VarScan v2.4.1 (Koboldt *et al*., 2012).

### Data analysis

All SNPs, in VCF format, were read into R using the `vcf2pooldata` function from the `poolfstat` package (Hivert *et al*., 2018; Gautier *et al*., 2022). The SNP data were filtered to include only SNPs with a minimum coverage of 60 reads, corresponding to 1x per haplotype in the pool. We further required the coverage of all pools to fall within the 0.1 to 99.9 quantile coverage thresholds and to have a minimum allele frequency of 0.008. All further analyses were performed in R using the `poolfstat` package. Genetic differentiation measured as Wright’s fixation index, F_ST_, was calculated using methods outlined in (Hivert *et al*., 2018). The geographic distance between stands was estimated from their respective latitude and longitude coordinates using the Haversine formula, which calculates the shortest distance, also known as the ‘great-circle-distance’, between two points using the `distHaversine` function from the `geosphere` package in R. To assess the genetic similarity between stands within a single locality, i.e. one old-growth and the two corresponding newly planted stands, we employed the outgroup *f*_3_ statistics (Patterson *et al*., 2012). Outgroup *f*_3_ statistics are a special case of how *f*_3_ statistics are commonly used. Instead of a target population and two possible source populations, as is used in calculating ordinary *f*_3_ statistics for admixture, the outgroup version uses two source populations and one outgroup (Patterson *et al*., 2012; Gautier *et al*., 2022). The test estimates the genetic similarity, also known as ‘shared genetic drift’^..^, between the source populations relative to the outgroup (Gautier *et al*., 2022). We calculated outgroup *f*_3_ tests using all combinations of old and planted stands within localities by cycling through which stand type was assigned as the outgroup. In this type of analysis, higher *f*_3_ statistics indicate that the two source populations are more genetically similar relative to the outgroup.

### Current and future climate data

Climate data for all stands were obtained from the geographic coordinates assigned to each stand during sampling using the ENVIREM database (http://envirem.github.io/, (Title & Bemmels, 2018) with a spatial resolution of 2.5 arcminutes (∼5 km). Climate data were obtained for either the current climate (1960-1990) or for the predicted climate in the year 2070 based on a representative future greenhouse gas concentration pathway (RCP4.5, (Moss *et al*., 2008). The RCP4.5 data were constructed based on climate data from Worldclim (v1.4, (Hijmans *et al*., 2005), http://worldclim.org/current), also at a 2.5 arcminute resolution, and converted to ENVIREM data format as previously described (Ingvarsson & Bernhardsson, 2020). Due to high collinearity among variables in the ENVIREM data, we selected seven representative climate variables that describe the variation in temperature and aridity across the study region (Supplementary Figure 1). The climate variables used in all further analyses are listed in Table 1 (Title & Bemmels, 2018).

**Table 1.** Climate variables and abbreviations used.

| Variable | Abbreviation | Brief description |
| --- | --- | --- |
| Thornthwaite aridity index | AIT | Index of the degree of water deficit below water need |
| Climatic moisture index | CMT | A metric of relative wetness and aridity |
| Continentality | CONT | Average temperature of warmest month - average temperature of coldest month |
| growingDegDays0 | DegD0 | Sum of mean monthly temperature for months with mean temperature greater than 0°C multiplied by number of days |
| growingDegDays5 | DegD5 | Sum of mean monthly temperature for months with mean temperature greater than 5°C multiplied by number of days |
| maxTempColdestMonth | MTC | The maximum temperature of the coldest month |
| PETColdestQuarter | PCQ | Mean monthly potential evapotranspiration (PET) of coldest quarter |

### Gene-environment association analyses

SNPs significantly associated with climate variables were identified in two ways. First, we selected SNPs based on the correlation between allele frequencies and the climate variable of interest across stands using the approach outlined by Fischer et al. (2013). Briefly, we generated 1,000,000 random allele frequency variables and associated each draw with one randomly selected climate variable to generate a null distribution for the correlation coefficient between allele frequencies and climate variables. We then used the 99.9% quantile of the resulting 1,000,000 correlation coefficients (i.e. *r*=0.730) as our threshold for selecting SNPs that were deemed to be significantly associated with a particular climate variable. If an SNP-environment correlation exceeded this threshold, the SNP locus was considered significantly associated with the environmental variable. We selected associated SNPs based on whether they were associated with a climate variable in old or planted stands separately. The second approach to identify outlier SNPs used latent factor mixed models (LFMM) (Frichot *et al*., 2013; Caye *et al*., 2019) as implemented in the `lfmm` package in R (https://CRAN.R-project.org/package=lfmm). The main benefit of the LFMM method is that it can adjust gene-environment association for the underlying population structure, something that the simple correlation method can not do. We first used a principal component analysis to estimate population structure in the SNP data (Supplementary Figure S2A), and based on the screeplot (Supplementary Figure S2B), we selected four (K=4) latent factors to correct for population structure in the data. In line with how we performed the correlation tests, we separately selected associated SNPs for old and planted stands.

To test for isolation by environment (Wang & Bradburd, 2014) we used partial Mantel tests to test for overall associations between genetic differentiation among stands and climate distance. This method has successfully been used in previous studies assessing associations between population differentiation and climate variables (e.g. Hancock *et al*., 2011; Nosil *et al*., 2012; Fischer *et al*., 2013). The method allows for comparing two pairwise distance matrices while controlling for the effect of a third. The dependent variable in all analyses was the pairwise F_ST_ matrix calculated based on outlier SNPs, selected based on either correlations or LFMM as outlined above. The predictor variable was the ‘environmental distance’ between sites, calculated by the Euclidean distances between sites for the corresponding environment variable using the `vegdist` function from the `vegan` package in R. The matrix used to control for the background population structure was a matrix with pairwise F_ST_ values calculated from all remaining SNPs after removing outlier SNPs. Partial Mantel tests were run using Pearson’s *r*, thus assuming a linear relationship between allele frequencies and environmental factors. The partial Mantel tests were implemented using the `mantel.partial` function from the `vegan` package in R.

Finally, we calculated the risk of non-adaptedness (RONA) (Rellstab *et al*., 2016) using all associated SNPs for the old and planted stands separately. RONA estimates the average expected allele frequency shifts required under future environmental conditions across all associated SNPs, with each locus weighted by the R^2^ value of the allele frequency-environment correlation (Pina-Martins *et al*., 2019). For the RONA analyses, we contrasted the current climate with the expected climate from the RCP4.5 greenhouse gas concentration scenario (Moss *et al*., 2008).

## Results

We generated over 1100 Gbp of sequencing data across the 45 sequence pools, with a median coverage of 1,850x per site across the 1.1Mb of the Norway spruce genome that our GBS analysis targeted (Supplementary Table S2). After SNP calling and filtering, 47,552 SNPs were retained and used in all subsequent analyses.

We used all SNP data from the complete set of pools of individuals to assess population structure in our collection of stands (Supplementary Figure S2). We also calculated pairwise F_ST_ values between all stands. Using a Mantel test, we assessed the evidence for isolation by distance by calculating the correlation between pairwise population differentiation and the great-circle distance between stands. Interestingly, we only detected weak but significant isolation by distance for the planted stands (old: *r*=0.075, *p*=0.254, planted: *r*=0.127, p=0.036, Supplementary Figure S3). When analysing different estimates of genetic diversity on a per-stand basis, we did not observe any significant differences between old and planted stands for the number of invariant sites, the proportion of rare alleles (alleles with a frequency <5%) or heterozygosity (Supplementary Figure S4). Finally, we assessed pairwise genetic differentiation (F_ST_) and genetic similarity (outgroup *f*_3_) between the different types of stands within each sample location. We observed a higher genetic differentiation between planted and old stands (Figure 2A) and a greater genetic similarity among the planted stands (Figure 2B).

**Figure 2.**
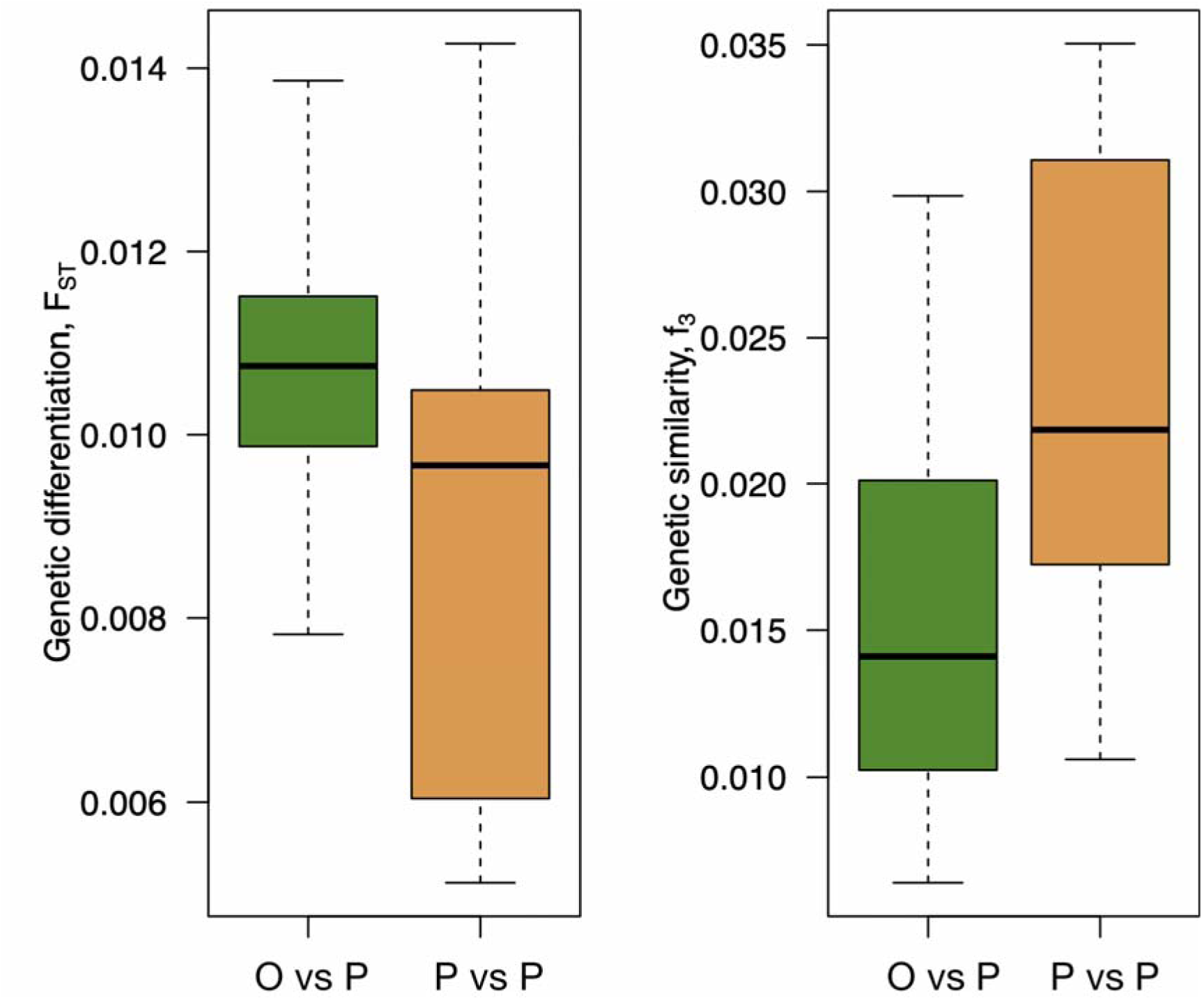
A) Pairwise genetic differentiation and B) pairwise genetic similarity (shared genetic drift, f_3_) between old and planted (O vs P) or between planted populations (P vs P) within each location.

We observe substantially greater numbers of significant associations between genetic diversity and climate in the old-growth stands when relying on simple allele-frequency correlations with the climate variables (the average number of significant correlations for old-growth and planted stands is 87 and 4, respectively, Table 2). We also observed strong, positive partial Mantel correlations for the old-growth stands with all climate variables. At the same time, only AIT was significant for the planted stands (Mean partial Mantel *r*= 0.914 and *r*=0.028 for old-growth and planted stands, respectively, Table 2, Figure 3). These correlations remain strong for old-growth stands under the projected climate for the RCP45 2070 scenario (mean partial Mantel *r=*0.833), while no such correlations were observed in the planted stands (mean partial Mantel *r=*-0.011, Table 2, Figure 3).

**Figure 3.**
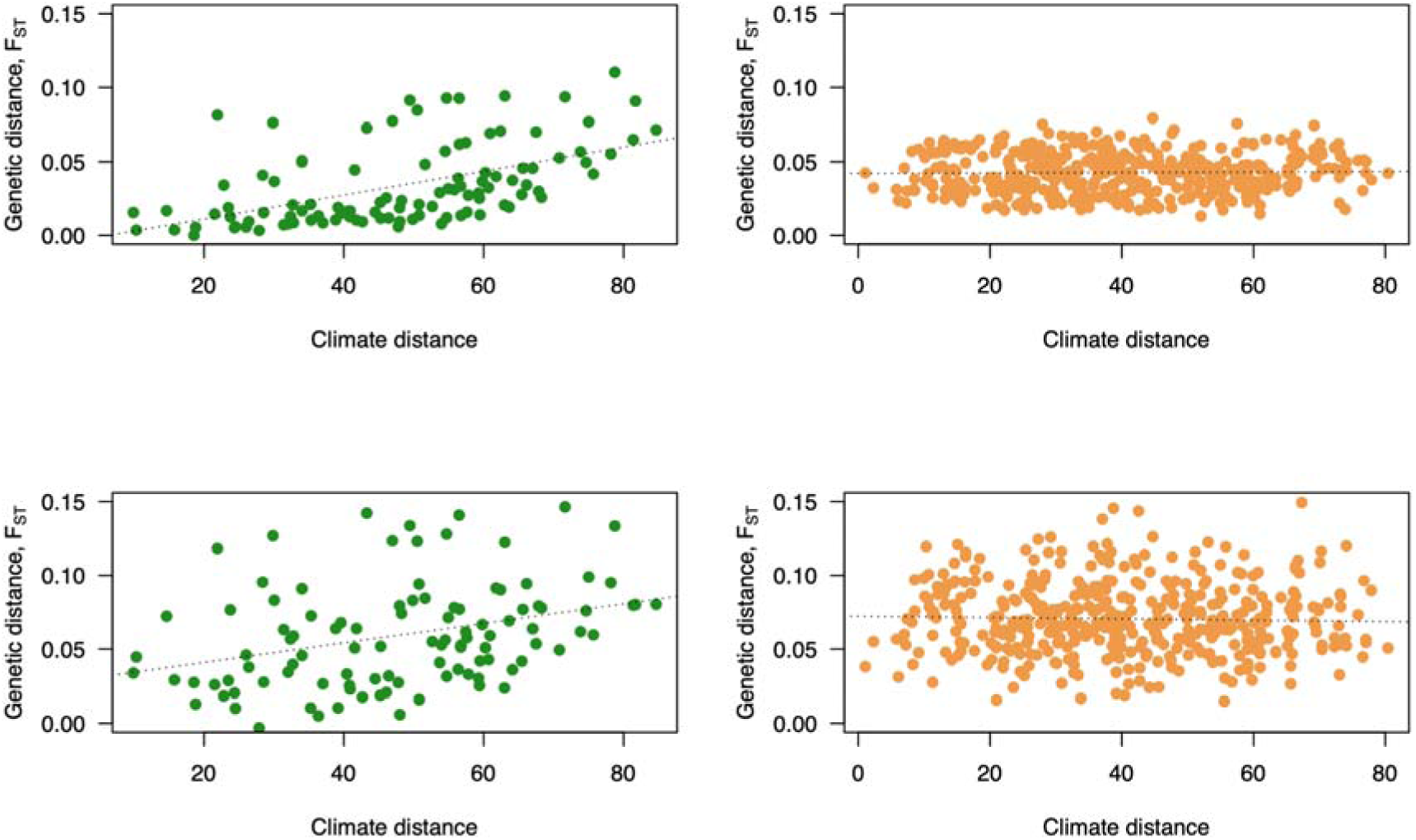
Associations between genetic and current climate distance for old and planted populations. In A) and B) the genetic distance between populations is calculated using all SNPs showing a significant correlation with one or more climate variables. In C) and D) SNPs significant in the LFMM analyses were used. In all comparisons, climate distance is calculated based on all seven climate variables.

**Table 2.**
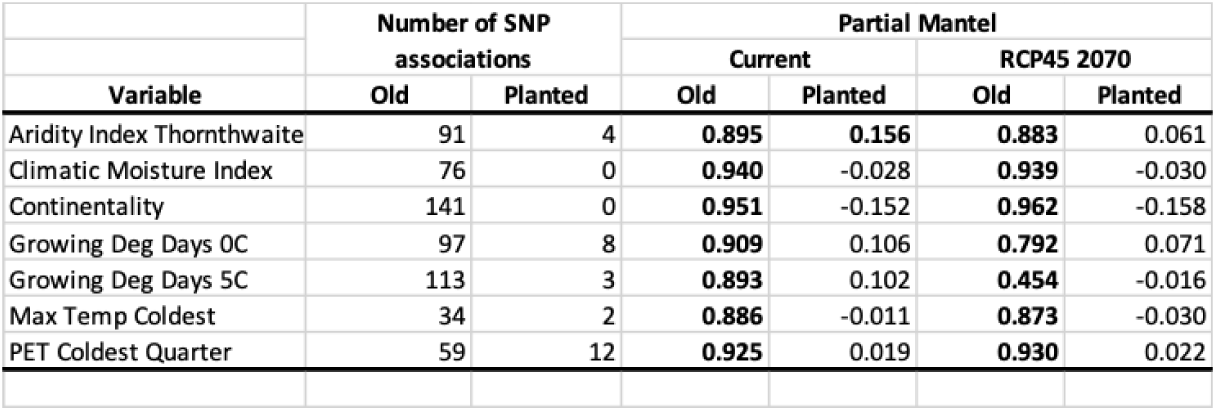
Associations and partial Mantel correlations for old-growth and planted stands using allele frequency correlations to select outlier loci.

We observed similar results when outlier SNPs were selected using LFMM, even if the overall patterns were not as strong as the analyses based on simple correlations. On average, 35 SNPs show significant associations with climate in old-growth stands compared to only 3 in the planted stands (Table 3) in the LFMM data set (Table 3). The partial Mantel correlations based on the LFMM-outlier SNPs for the old-growth stands were also high and significant for all climate variables (mean partial Mantel *r*=0.715). At the same time, only CONT was significant for the planted stands (mean partial Mantel *r*=0.016, Figure 3). For the 2070 RCP45 climate scenario, all climate variables, except DegD5, remained significantly correlated for old-growth stands (mean partial Mantel *r*=0.645, Table 3, Figure 3). For the planted stands, none of the partial Mantel correlations were significant for this climate scenario (mean partial Mantel *r*=0.008, Table 3, Figure 3).

**Table 3.**
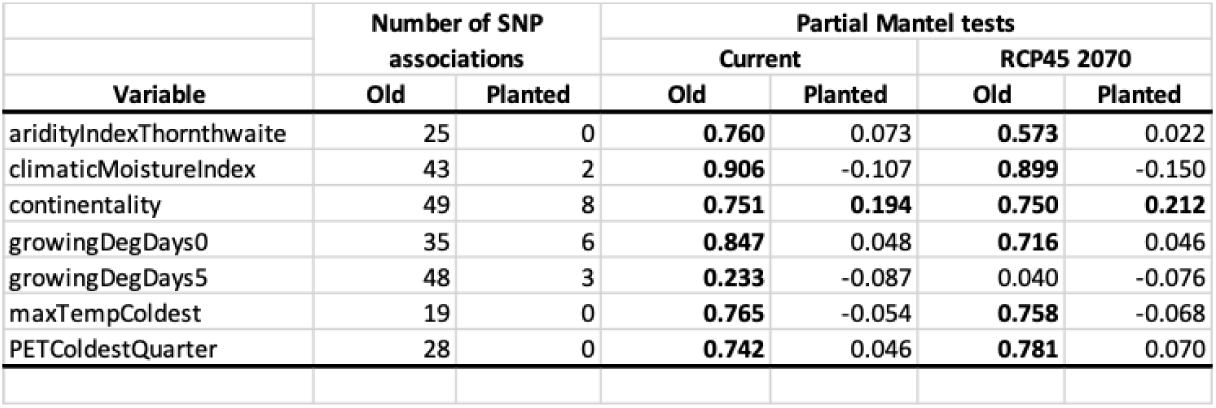
Associations and partial Mantel correlations for old-growth and planted stands using LFMM to select outlier loci.

The planted stands consistently display greater estimates of the risk of non-adaptedness (RONA) for all the different climate variables we assessed, regardless of whether the RONAs were estimated using outlier loci selected based on allele frequency correlations or using LFMM (Figure 4A and B). The only cases where RONA was not significantly greater in newly planted stands were DegD5, MTC and PCQ for the LFMM-based calculations (Figure 4B).

**Figure 4.**
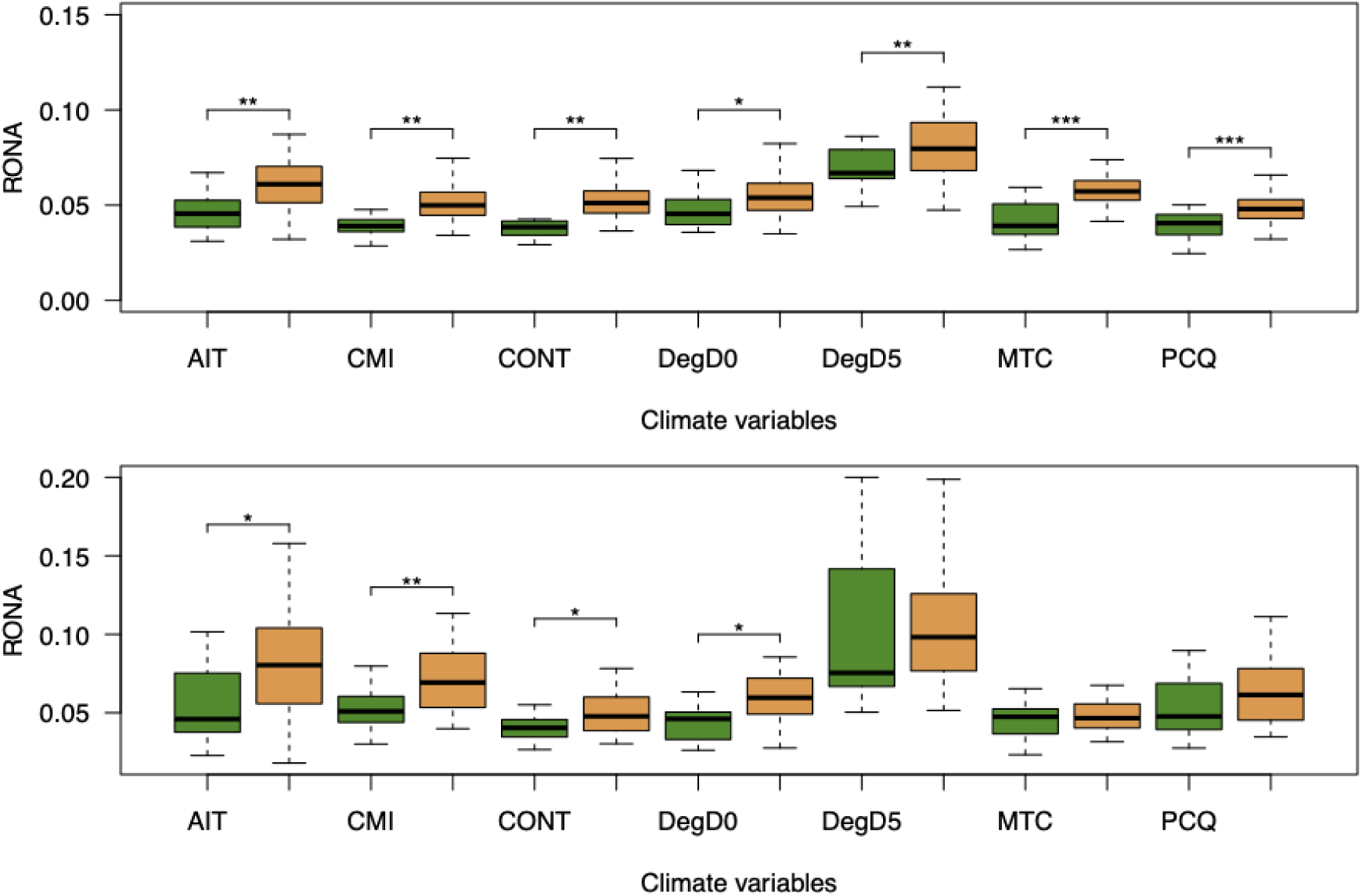
Risk of non-adaptedness (RONA) for old-growth (green) and newly planted (green) stands for all climate variables (see Table 1 for an explanation of abbreviations). In A), outlier loci are identified using allele frequency correlations, and in B), outlier loci are identified using LFMM.

## Discussion

We did not detect any systematic differences in the amount of genetic diversity maintained in old versus planted populations, regardless of what measure we used (Supplementary Figure 4). However, we observe greater genetic differentiation among old-growth versus planted stands and greater genetic similarities between the planted stands within sites (Figure 2). The lack of differences in estimates of genetic diversity between stand types is unsurprising since even relatively small samples, around 20-25 individuals, are expected to capture most of the diversity available in a source population (Ingvarsson & Dahlberg, 2019). Our results are also in line with earlier studies that have assessed genetic diversity in forest trees and which have generally found no or only minor differences in estimates of genetic diversity between natural and managed stands of forest trees (Bergmann & Ruetz, 1991; Maghuly *et al*., 2006; Garcia-Gíl *et al*., 2015; Sønstebø *et al*., 2018; Ruņģis *et al*., 2019). Previous studies have used several different types of genetic markers, ranging from allozymes to microsatellites or RAPDs, and they have mostly failed to detect significant differences in genetic diversity between planted and natural stands. Bergmann and Ruetz (1991) used eight enzyme loci, to compare 45 seed orchard trees with 60 random spruce trees from the same area and found a significant difference in average heterozygosity. Ruņģis et al. (Ruņģis *et al*., 2019) found higher numbers of alleles and higher average gene diversity in naturally regenerated forests (153 trees) in comparison to progeny from seed orchards (144 trees) when genotyping using 11 SSRs. Maghuly et al. (2006) also used SSRs, chloroplast SSRs, and mitochondrial markers to compare two age classes and three populations at different elevations. From 50 old and 100 young individuals from each population, they found no difference in seven chloroplast SSRs. Their five SSRs indicated that there was less genetic diversity and heterozygosity among the different populations than there was within populations. Sønstebø et al. (2018) compared two seed orchards (25 and 60 parents) with semi-natural forests and unmanaged stands. Four samplings were done with 300 seeds in each seed orchard, and 13 seed lots were collected from semi-natural forests. Finally, needles were sampled from five natural forests. With 11 microsatellites, they showed a slightly lower genetic diversity in the form of allelic richness from the seed orchard samples, with the most pronounced difference seen in the orchard with only 25 parent trees (Sønstebø *et al*., 2018). One thing these studies have in common is that they have all relied on a relatively modest number of independent genetic markers, suggesting that the power to detect any differences in genetic diversity has been low in these studies.

Compared to earlier studies, we used a substantially greater number of independent genetic markers in the current study (∼47k SNPs), and we also had reasonable coverage of the spruce genome (>8k unique genomic regions). We have also tried to sample from a relatively large geographic area in northern Sweden, spanning the Baltic coast to the Scandinavian mountains (∼25000 km^2^). We included 45 different stands and a total of 1350 trees in our study that, combined with a large number of markers, should give our study a substantially greater statistical power for detecting possible differences in genetic diversity and differentiation. This allows us to confidently state that we do not observe any differences in genetic diversity between old and planted populations. If such differences nevertheless exist, they must be small. The seeds used for re-planting trees at the planted stands in our study are most likely derived from the second round of tree breeding (established 1981-1994) and are not representative of the most advanced selection made to date. It is possible that further selection rounds would create larger differences in genetic diversity between natural and production forests and make such differences easier to detect, but this remains to be investigated.

It is, however, essential to distinguish between putatively neutral genetic variation and genetic variation linked explicitly to traits that confer adaptive traits. Most of the variation found across the genome of a species is likely neutral or nearly neutral, and only a small fraction of all variants are involved in mediating adaptive traits. We know from many earlier studies that the genetic structure of adaptive variation is often fundamentally different from neutral genetic variation (Le Corre & Kremer, 2003; Savolainen *et al*., 2013). Natural selection increases differentiation among populations for adaptive traits that contribute to local adaptation. Local adaptation persists in most forest trees despite high levels of gene flow that continuously introduces potentially maladaptive genetic variation (Kremer *et al*., 2012). As forest tree populations are nevertheless able to adapt to local environments in the face of extensive gene flow, natural selection acting on traits that confer local adaptation must be strong enough to overcome the homogenising effects of gene flow. So even if neutral markers display low genetic differentiation, consistent with extensive gene flow among populations, patterns of genetic differentiation at loci directly involved in controlling adaptive traits could be quite different, resulting in patterns of *isolation by environment* (IBE), where genetic and environmental distances are positively correlated, independent of their geographic distance (Wang & Bradburd, 2014).

In line with these observations, a strikingly different picture emerges when we focus on adaptive genetic variation in the old-growth and planted stands. The amount of variation and how this variation is associated with underlying climate variables drastically differ between the two types of forest stands we have studied (Figure 3, Tables 2 and 3). We observe strong and consistent correlations between genetic and environmental distance in the old-growth populations (Figure 3, Tables 2 and 3), and this holds regardless of the method used to identify outlier loci assumed to represent adaptive variation. In contrast, we observe no or only weak correlations when assessing the same loci in the planted stands (Figure 3, Tables 2 and 3). We also observe a ten to twenty-fold greater number of significant SNP-climate associations in the old-growth forests than in the planted populations, regardless of whether we use current or RCP45 climate variables (Tables 2 and 3). When we use these outlier SNPs to assess isolation by environment and control for the background genetic structure, we consistently observe strong, positive correlations in the old-growth forests. At the same time, this pattern is largely absent for the planted stands (Tables 2 and 3). Similarly, when assessing the risk of non-adaptedness (RONA) of current standing variation under a scenario of future climate warming (RCP45), we observe that RONA estimates are consistently and significantly greater for the planted populations, suggesting that they are at greater risk of suffering adverse effects due to climate warming.

These results highlight a possible underappreciated consequence of large-scale reforestation programs in that they may weaken local adaptation patterns in managed populations planted with material taken from non-local sources, such as seed orchards or seed collections from natural (but not necessarily local) populations. This suggests that planted populations could be exposed to greater risks from climate change if care is not taken to include local adaptation as a key criterion when selecting seed source material. Forest tree breeding is usually performed with local adaptation and future climate change in mind (Cortés *et al*., 2020), but this is often done using proxies for climate change, e.g moving material across latitudinal gradients. The risk with this approach is that it fails to account for more fine-grained patterns of local adaptation or scenarios where local adaptation is driven by non-clinal allele frequencies (Lotterhos, 2023). The old-growth populations we have assessed show strong associations with local climate and may thus harbour important adaptive alleles that, for various reasons, are either absent or occur at low frequencies in source populations used for reforestation of the planted stands. The old-growth populations may thus serve as important sources for such adaptive alleles, and future work should focus on identifying these alleles and making a dedicated effort to introduce them into breeding populations and relevant seed orchards used in reforestation.

An important caveat of climate vulnerability estimates based on allele frequency differences across environments is that they only serve as proxies for potential adaptive mismatches rather than direct assessments of fitness (Lotterhos, 2024; Lind *et al*., 2024). These estimates rely on the degree of genetic change needed for a population to adapt to future conditions. Still, they do not directly measure the actual reproductive success, survival, or overall fitness of individuals in a changing environment (Lotterhos, 2024; Lind *et al*., 2024; Lind & Lotterhos, 2024). While allele-frequency-based estimates correlate with fitness declines in controlled experiments such as common garden studies, they do not inherently account for other ecological factors influencing fitness in natural settings, making them proxies rather than direct measures of maladaptation or vulnerability. Forest tree breeding programs often use long-term provenance trials where genetic material from various populations (provenances) is tested across different environments to identify genotypes with broad climatic adaptability (Cortés *et al*., 2020). So, rather than focusing on matching local adaptation at target planting sites, forest tree breeding often uses multi-trait selection indices to balance productivity with resilience traits and emphasise genetic diversity to buffer against unpredictable stresses (Cortés *et al*., 2020).

## Conclusions

This study provides a comprehensive comparison of genetic diversity and adaptive potential between old-growth and recently planted stands of Norway spruce (*Picea abies*) in northern Sweden. Utilising extensive genomic data and broad geographic sampling, our results reveal no significant differences in neutral genetic diversity between natural and planted populations, suggesting that current forestry practices, including seed orchard selections and reforestation methods, have not substantially reduced overall genetic variation. However, the patterns are strikingly different when assessing adaptive variation, which is crucial for predicting future responses to climate change. Old-growth forests show substantially stronger patterns of local adaptation and are also predicted to suffer lower risks due to future climate change. These findings underscore the importance of maintaining and enhancing adaptive genetic diversity through forest management to ensure resilience under future climate change. While planting genetically improved seedlings from seed orchards does not diminish overall genetic diversity in Norway spruce populations and helps accelerate reforestation efforts, careful consideration is necessary to ensure that efforts are taken to preserve genetic variation essential for local adaptation. Old-growth forests could serve as important sources for (re-)introducing such adaptive variation into existing tree breeding programs.

## Data availability

The GBS raw reads have been deposited in NCBI’s Sequence Read Archive (SRA) under accession number PRJEB89643 (https://www.ncbi.nlm.nih.gov/bioproject/PRJEB89643/). Background information on all sites, environmental data at the site of origin for all populations and the PoolSeq SNP data are available from Zendo (https://doi.org/10.5281/zenodo.XXXX) under a CC BY-SA 4.0 license. All scripts used for the analyses described in the paper are available on GitHub under a MIT License (https://github.com/parkingvarsson/SpruceOldvsPlanted).

## Author contributions

HE and PKI conceived of and designed the experiments. HE did the field sampling and prepared GBS libraries. CB handled the GBS sequencing data and performed the read mapping. PKI performed the PoolSeq SNP calling. HE and PKI carried out all population genetic analyses. HE and PKI wrote the paper. All authors commented on the manuscript. All authors have read and approved the final version of the manuscript.

## Acknowledgements

The research has been funded by grants from the Swedish Foundation for Strategic Research (SSF Grant No. RBP14-0040) and the Knut and Alice Wallenberg Foundation. All analyses were performed using resources provided by the National Academic Infrastructure for Supercomputing in Sweden (NAISS), partially funded by the Swedish Research Council through grant agreement no. 2022-06725, through the Uppsala Multidisciplinary Centre for Advanced Computational Science (UPPMAX) under the compute projects SNIC 2017/1-438, SNIC 2018/3-529, SNIC 2019/3-555 and the storage projects uppstore2017066 and uppstore2017145.

