## Supplementary Tables and Figures for "Contrasting patterns of local adaptation and climate resilience across forest management regimes in Norway spruce (*Picea abies):* implications for reforestation practices under climate change"

**Supplementary Table S1**. Type, location and age of sampling sites and summary statistics of sequencing data per library.

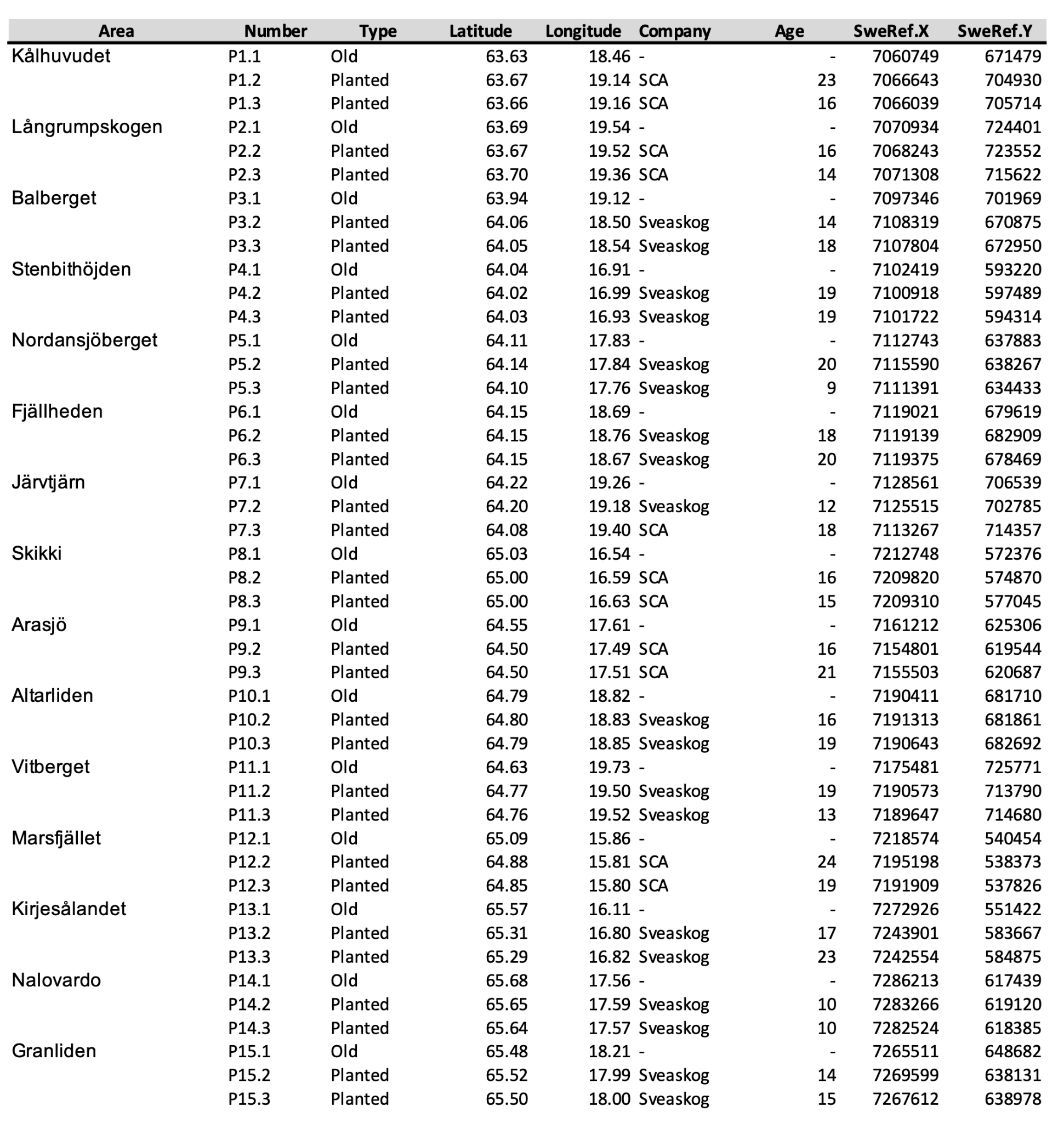

**Supplementary Table S2.** Summary statistics of the sequencing data.

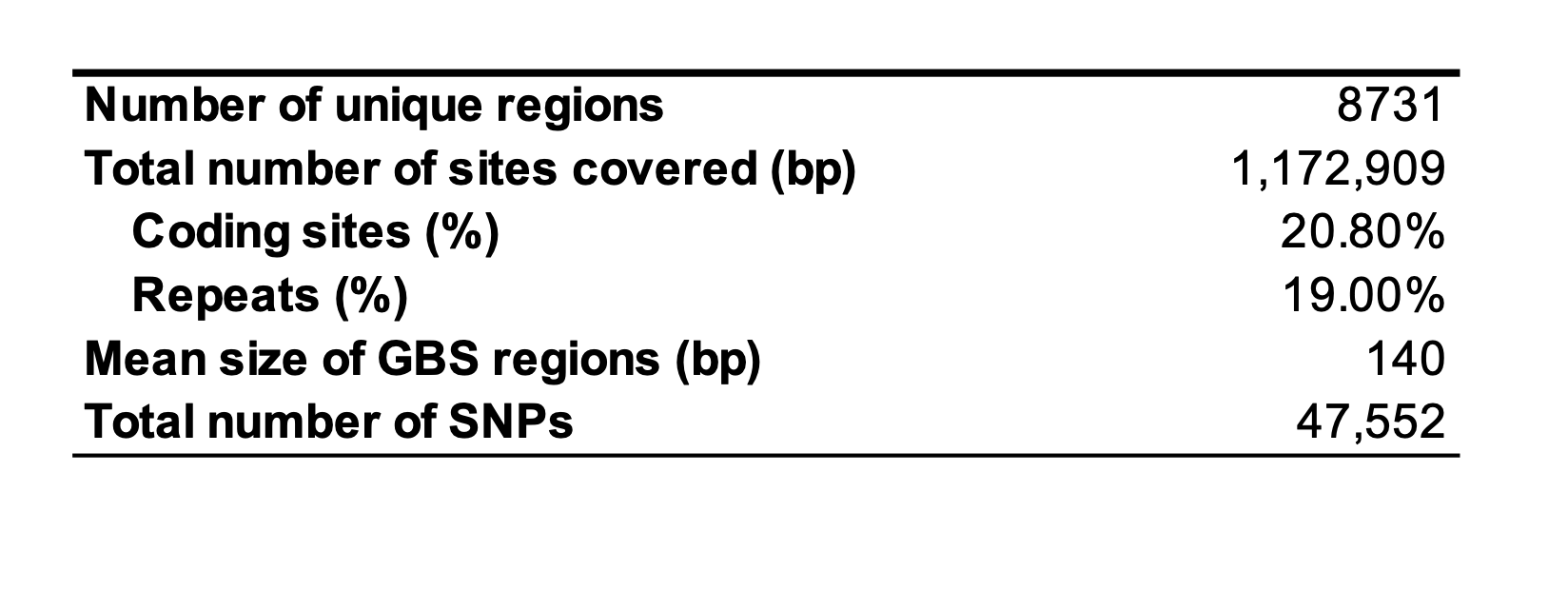

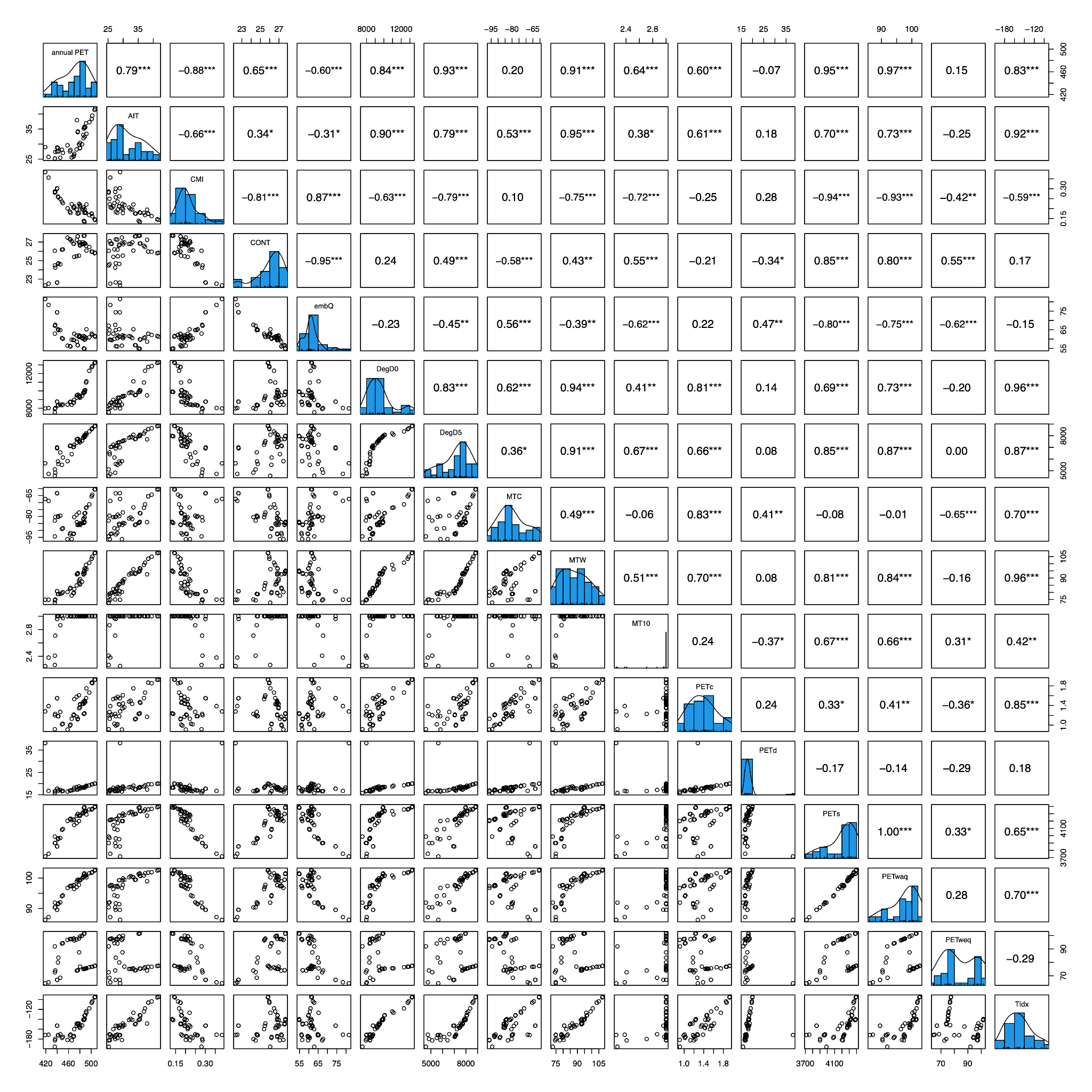
**Supplementary Figure S1.** Summary of the ENVIREM climate variables for all populations used in the study. The diagonal displays the distribution of the individual variables, figures below the diagonal display pairwise scatterplots for all variables and above the diagonal, the corresponding correlation coefficients are given. **p*<0.05, ***p*<0.01, ****p*<0.001

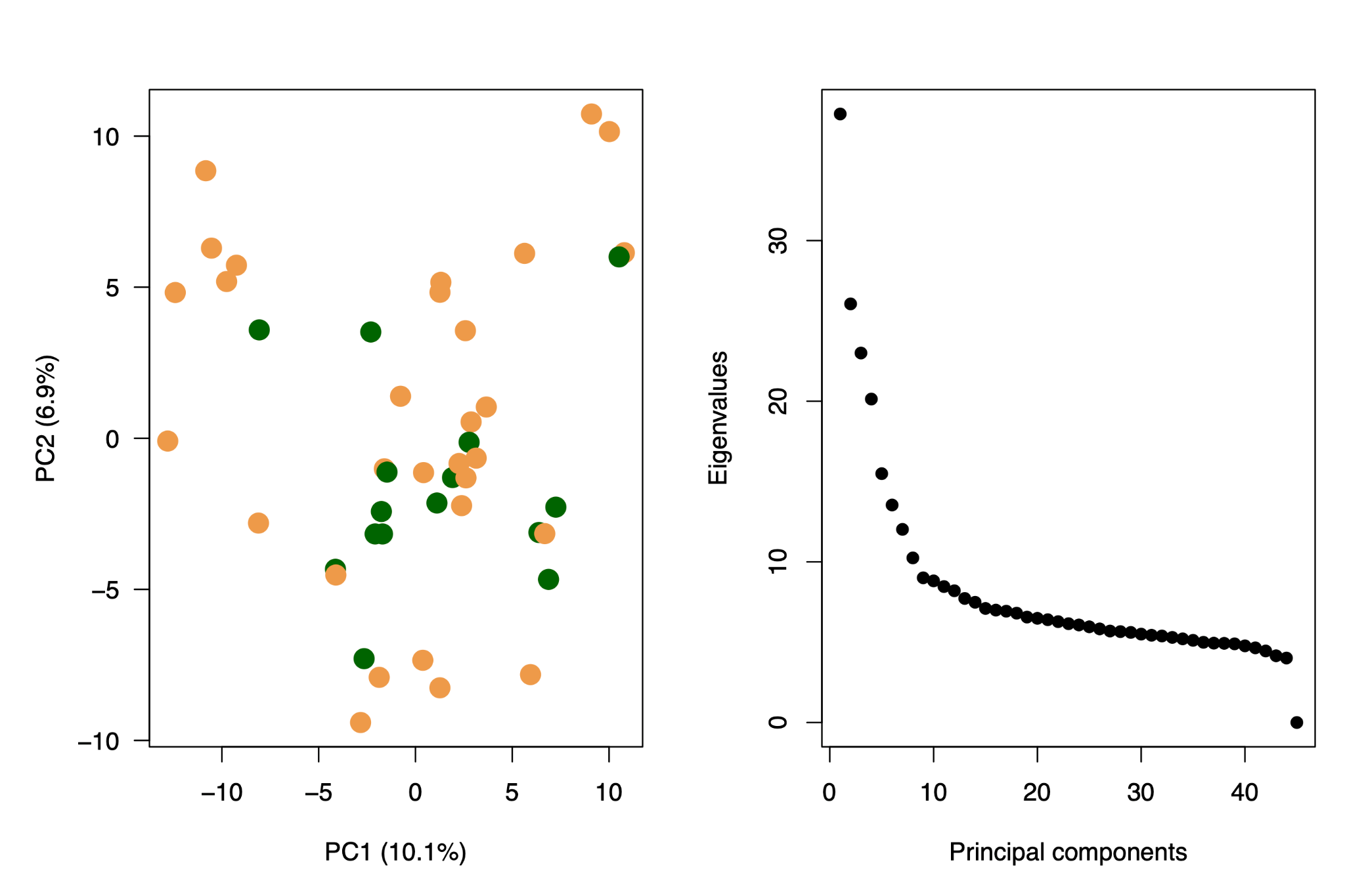

**Supplementary Figure S2.** A) Principal component analysis and B) screeplot of the allele frequency data. Old and planted populations in A) are depicted using green and tan, respectively.

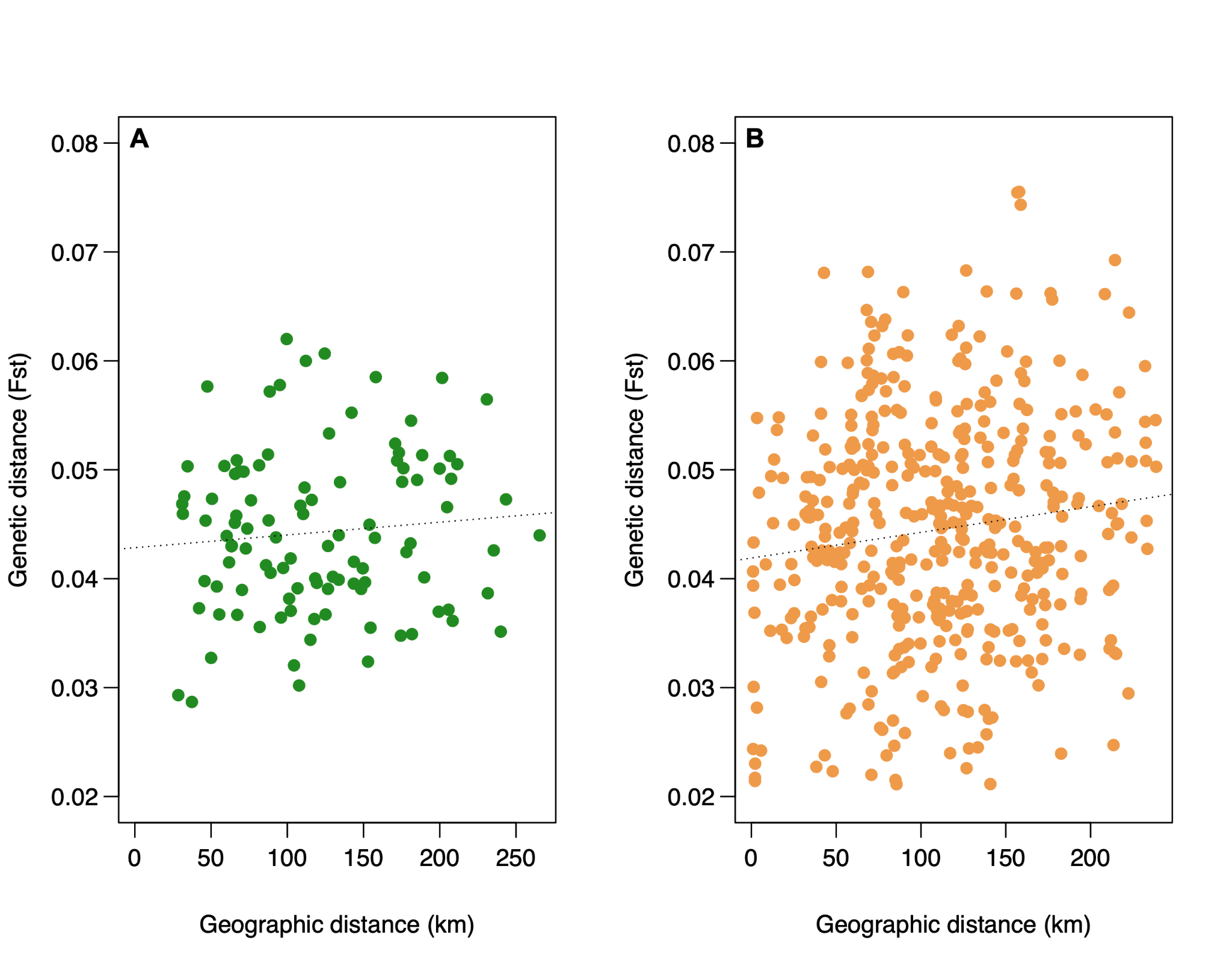

**Supplementary Figure S3.** Isolation by distance for A) old and B) planted populations. The dotted line is the best-fitting regression line and is only used for visual guidance.

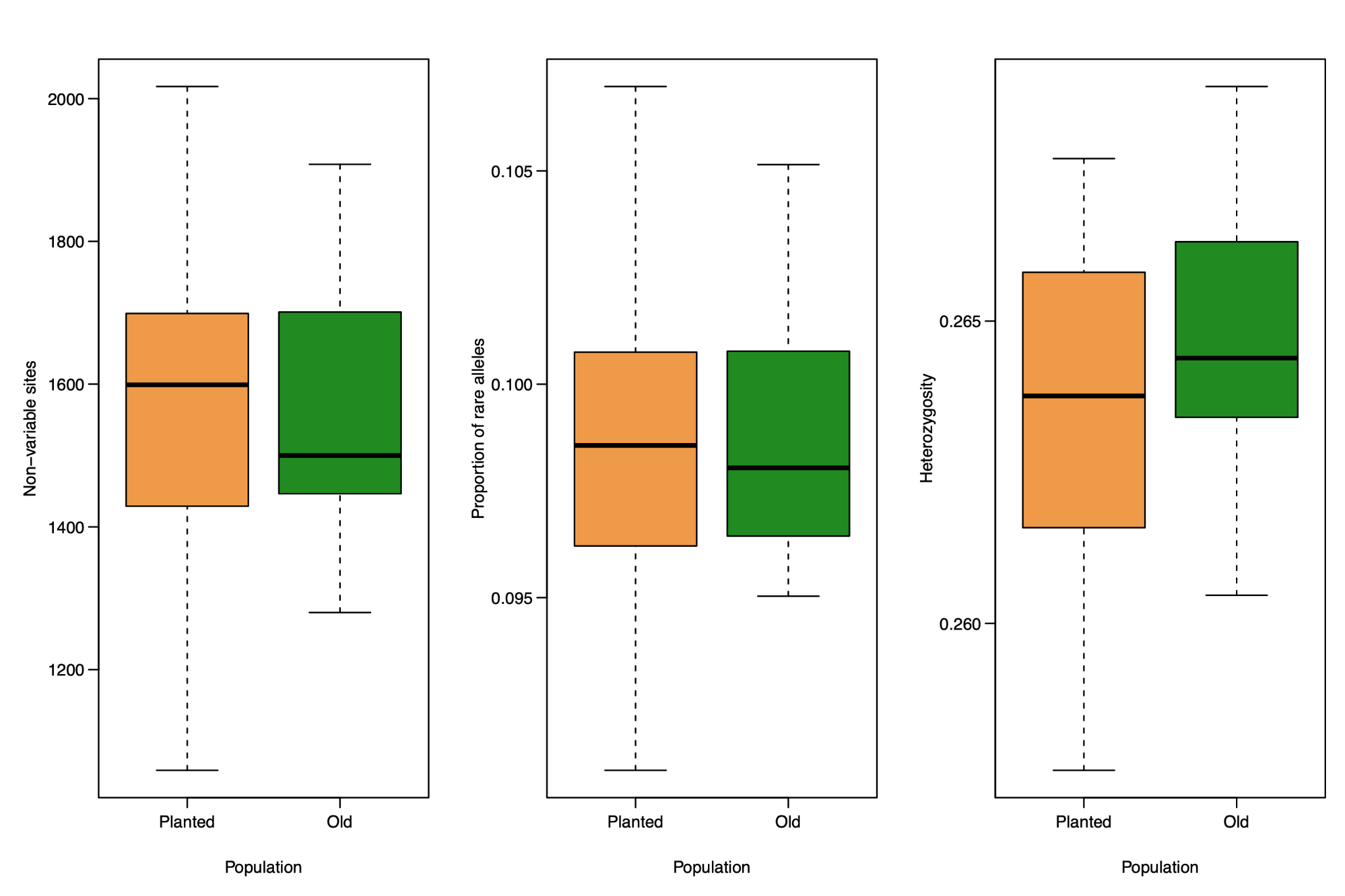

**Supplementary Figure S4.** The number of invariable sites, B) the proportion of rare alleles (alleles with frequency <0.05) and C) heterozygosity in planted (tan) and old (green) populations. None of the population comparisons are significantly different.
